# Age-related changes in the motor planning strategy slow down motor initiation in elderly adults

**DOI:** 10.1101/2020.05.18.101774

**Authors:** Nikita S. Frolov, Elena N. Pitsik, Vladimir A. Maksimenko, Vadim V. Grubov, Anton R. Kiselev, Zhen Wang, Alexander E. Hramov

**Author notes:** These authors contributed equally to this work (Nikita S. Frolov, Elena N. Pitsik, Vladimir A. Maksimenko, Alexander E. Hramov).

## Abstract

Age-related changes in the human brain functioning crucially affect the motor system, causing increased reaction time, low ability to control and execute movements, difficulties in learning new motor skills. The lifestyle and lowered daily activity of elderly adults, along with the deficit of motor and cognitive brain functions, might lead to the developed ambidexterity, i.e. the loss of dominant limb advances. Despite the broad knowledge about the changes in cortical activity directly related to the motor execution, less is known about age-related differences in the motor initiation phase. We hypothesize that the latter strongly influences the behavioral characteristics, such as reaction time, the accuracy of motor performance, etc. Here, we compare the neuronal processes underlying the motor planning of fine motor tasks between elderly and young subjects. We demonstrate that aging significantly reduces the speed of motor initiation in the dominant hand task due to the different motor planning strategies employed by elderly and young adults. Based on the results of the whole-scalp electroencephalography (EEG) analysis, we suggest that young adults tend to use the efficient and fast mechanism of motor working memory. In contrast, elderly adults involve a more demanding sensorimotor integration process similar to the non-dominant hand task.

## Introduction

Healthy aging affects neural processes by changing the neurochemical and structural properties of the brain [1]. It determines the cognitive and motor performance decline during a daily activity of elderly adults and negatively influences the quality of their life. The markers of age-related neural impairments are observed at the behavioral level as slowing of the reaction time (RT), reduced motor control and coordination, etc. [2, 3].

Upper limbs represent the most active part of the human motor system; thus, the degradation of its functioning with age is the most prominent [4]. Plenty of studies report difficulties in accomplishing complex motor tasks related to the deficit of hand movement coordination, ability to control force, execute sequential actions, learn new motor skills, etc. [2, 5, 6]. However, the motor performance decline while executing fine motor tasks is also well-documented [7–9]. Several studies relate these phenomena to the over-activation of the brain motor and prefrontal area, which controls the motor execution process [10–13]. Specifically, the recruitment of additional ipsilateral motor regions in elderly adults is supposed to provide a compensatory mechanism that supports overcoming the age-related structural changes in the human brain [14, 15]. On the one hand, it helps to maintain the performance of executed motor actions. On the other hand, this mechanism demands more neuronal resources and, therefore, slowers the motor response. Also, several studies relate the over-activation of cortical areas to the ‘use-dependent plasticity’ [16], which is supposed to underly dedifferentiation of brain functions in advanced age. In the context of the motor system, it is manifested as developed ambidexterity, i.e., the loss of the dominant limb advances [8, 17].

While the age-related differences cortical activation directly related to motor execution and control is extensively studied, less is known about the effect of healthy aging on the motor planning phase and its influence on RT. Exploring these mechanisms is crucial to deeper understand motor control in humans. Motor planning is also subjected to the age-related changes due to the following: (i) motor initiation process involves many higher cognitive functions such as sensory processing, working memory, motor embodiment, and sensorimotor integration [18–21], which are known to decline strongly with age; (ii) the theta activity underlying the majority of these processes exhibits significant age-related changes – abnormally increased theta activity in elderly people indicates subjective cognitive dysfunction and suspected dementia [22, 23].

Based on the above, we hypothesize that the age-related changes in the motor planning mechanism also affect the slowing of the motor initiation phase in elderly adults. To address the issue, we considered the differences in cortical activity during the controlled execution of fine motor tasks between elderly adults and young adults using electroencephalography (EEG). Consistent with the dedifferentiation theory [8, 17], we found that the motor cortex of younger adults activated much faster during the dominant hand task, while in elderly adults, the time required for motor activation was equal for both hands and approached the level of the non-dominant hand of younger adults. Further, as expected, we found significant differences in cortical activation during the time interval preceding the motor action. In elderly adults, as well as in young adults performing the non-dominant hand task, we observed the increased theta-band power in sensorimotor and frontal areas, whereas theta-activation was insignificant in young adults during the dominant hand task. Finally, based on the results of between-subject functional connectivity analysis, we revealed that motor planning involves different types of cortical interactions in young adults and elderly adults, which allows concluding about age-related changes in motor planning mechanisms.

## Materials and methods

### Participants

Two groups of healthy volunteers, including 10 elderly adult subjects (EA group; age: 65±5.69 (MEAN±SD); range: 55-72; 4 males, 6 females) and 10 young adult subjects (YA group; age: 26.1±5.15 (MEAN±SD); range: 19-33; 7 males, 3 females), participated in this study. All subjects were right-handed and had no history of brain tumors, trauma or stroke-related medical conditions. The experimental protocol was approved by the local research Ethics Committee of Innopolis University. The experimental study was performed in accordance with the Declaration of Helsinki. All participants were pre-informed about the goals and design of the experiment and signed a written informed consent.

### Task

All participants were instructed to sit on the chair with their hands lying comfortably on the table desk in front of them, palms up. The timeline of the experimental session is presented in Fig. 1A. First, the background EEG was recorded at the beginning of the experiment when the participants were instructed to sit relaxed, eyes open, and not to think about anything particular (5 minutes). The active phase of the experiment included sequential execution of 60 fine motor tasks. Each task required squeezing one of the hands into a wrist after the audio signal and holding it until the second signal (30 tasks per hand). The duration of the signal determined the type of movement: short beep (0.3 s) was given to perform a non-dominant hand (left hand, LH) movement and long beep (0.75 s) was given to perform a dominant hand (right hand, RH) movement. Thus, we conducted a mixed-design experimental study with the Movement Type (LH and RH conditions) as within-subject factor and the Age (EA and YA groups) as between-subject factor.

**Fig 1.**
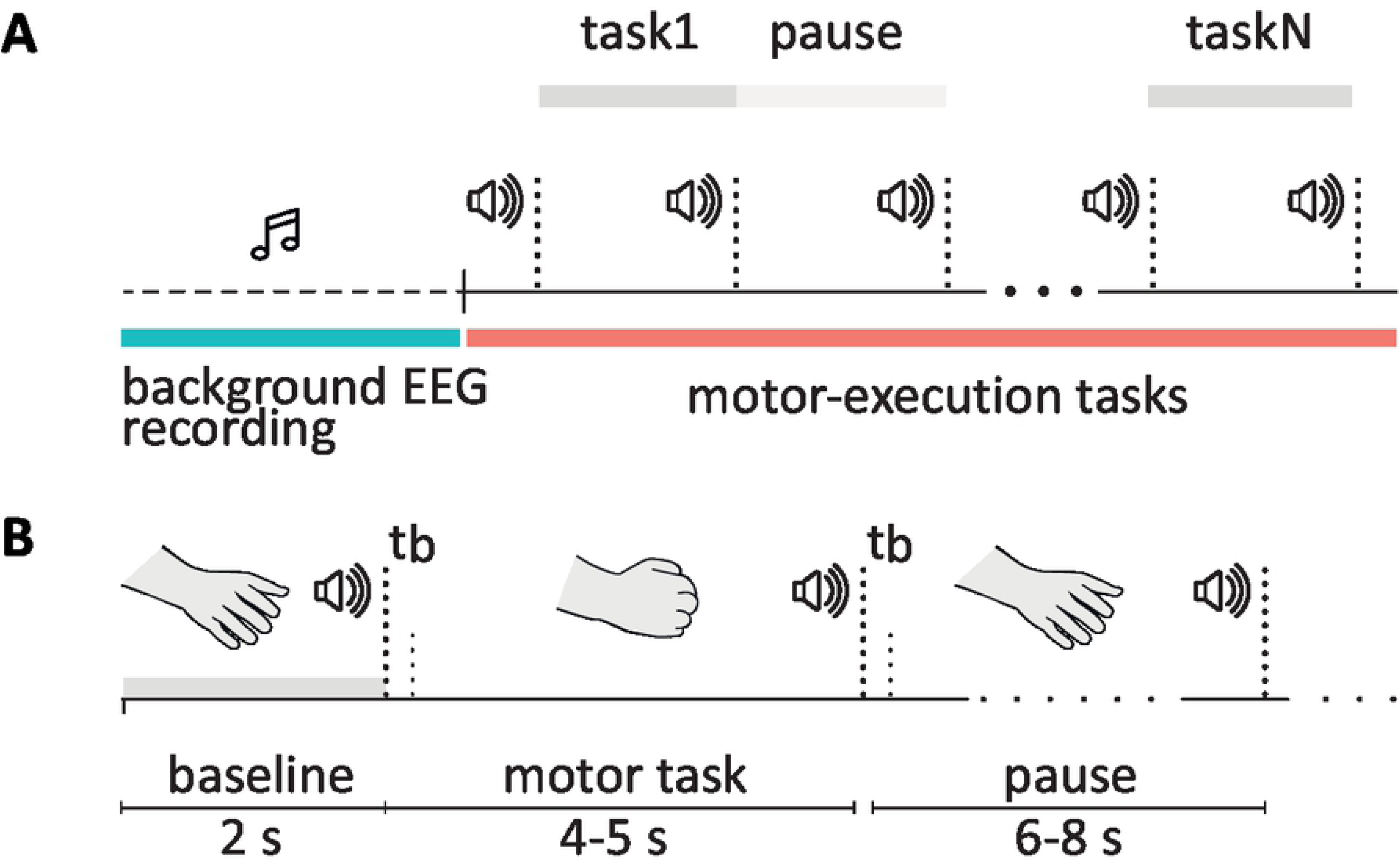
Experimental paradigm. Timelines of the experimental session (A) and a single motor task (B). Here, *t*_*b*_ is the duration of the beep, which is 0.3 s for the LH movement command, and 0.75 s for the RH movement command.

The timeline of a single task movement is presented in Fig. 1B. The time interval between the signals during the task and the pause between the tasks were chosen randomly in the range 4–5 s and 6–8 s, respectively. The types of tasks were mixed in the course of the session and given randomly to exclude possible training or motor-preparation effects caused by the sequential execution of the same tasks. The overall experimental session lasted approximately 16 minutes, including the background cortical activity recording and series of motor task execution.

### EEG data acquisition and preprocessing

We acquired EEG signals using the monopolar registration method (a 10—10 system proposed by the American Electroencephalographic Society [24]). According to this, we recorded EEG signals with 31 sensors (O2, O1, P4, P3, C4, C3, F4, F3, Fp2, Fp1, P8, P7, T8, T7, F8, F7, Oz, Pz, Cz, Fz, Fpz, FT7, FC3, FCz, FC4, FT8, TP7, CP3, CPz, CP4, TP8) and two reference electrodes A1 and A2 on the earlobes and a ground electrode N just above the forehead. We used the cup adhesive Ag/AgCl electrodes placed on the “Tien–20” paste (Weaver and Company, Colorado, USA). Immediately before the experiments started, we performed all necessary procedures to increase skin conductivity and reduce its resistance using the abrasive “NuPrep” gel (Weaver and Company, Colorado, USA). We controlled the variation of impedance within a range of 2–5 kΩ during the experiment. The electroencephalograph “Encephalan-EEG-19/26” (Medicom MTD company, Taganrog, Russian Federation) with multiple EEG and two EMG channels performed amplification and analog-to-digital conversion of the recorded signals. The EMG signals were acquired to verify the correctness of the epochs segmentation. This device possessed the registration certificate of the Federal Service for Supervision in Health Care No. FCP 2007/00124 of 07.11.2014 and the European Certificate CE 538571 of the British Standards Institute (BSI).

The raw EEG and EMG signals were sampled at 250 Hz and filtered by a 50–Hz notch filter by embedded hardware-software data acquisition complex. Additionally, raw EEG signals were filtered by the 5th-order Butterworth filter with cut-off points at 1 Hz and 100 Hz. Eyes blinking and heartbeat artifact removal was performed by the Independent Component Analysis (ICA) [25]. The recorded EEG and EMG signals presented in proper physical units (millivolts) were segmented into four sets of epochs according to the (group [YA, EA], condition [LH, RH]) combinations: YA LH, YA RH, EA LH, and EA RH. Each epoch was 10 s long, including 2s baseline activity and 8s motor-related activity. Data was then inspected manually and corrected for remaining artifacts. Epochs which we failed to correct manually mostly due to the strong muscle artifacts were rejected. Finally, each set contained 15 corrected epochs, which was equal to the minimal number of the artifact-free epochs over all participants.

All preprocessing steps including filtering, artifact removal and epoching were performed using MNE package (ver. 0.20.0) for Python 3.7 [26]. The analyzed EEG data is available online [27].

### Time-frequency analysis in sensor space

For each (group, condition)–set of epochs, we estimated spectral power in theta (4-8 Hz) and alpha/mu (8-14 Hz) frequency bands using time-frequency analysis implemented in MNE. Particularly, time-frequency representation of the EEG epochs was obtained via Morlet complex-valued wavelets in the range 4-30 Hz and contrasted with 2s baseline period using ‘percent’ mode, i.e., subtracting the mean of baseline values followed by dividing by the mean of baseline values. The number of cycles in the wavelet transform was set for each frequency *f* as *f/*2. Then, obtained time-frequency representations were averaged over epochs for each subject.

#### Estimation of the motor brain response time

A priory knowledge about the cortical activation during movements execution implies that motor brain response is determined as a pronounced event-related desynchronization (ERD) of mu-oscillations in the contralateral area of the motor cortex. Notably, a wide body of EEG studies reports that symmetrical C4 and C3 sensors evidence brain motor response in case of the left- and right-hand movements, respectively [28–31]. Here, we used mu-band event-related spectral power (ERSP_*µ*_) at C4 and C3 sensors to estimate motor brain response time (MBRT) associated with LH and RH conditions for each subject of both groups. We manually inspected each time-series and defined MBRT as the first minimum of the mu-band spectral power below the 2.5th baseline level (Fig. 2, A). Thus, we collected four sets of MBRT corresponding to each (group, condition)-set. Statistical comparison of the MBRT was performed using a two-way mixed-design ANOVA test implemented in JASP open-source statistical software [32].

**Fig 2.**
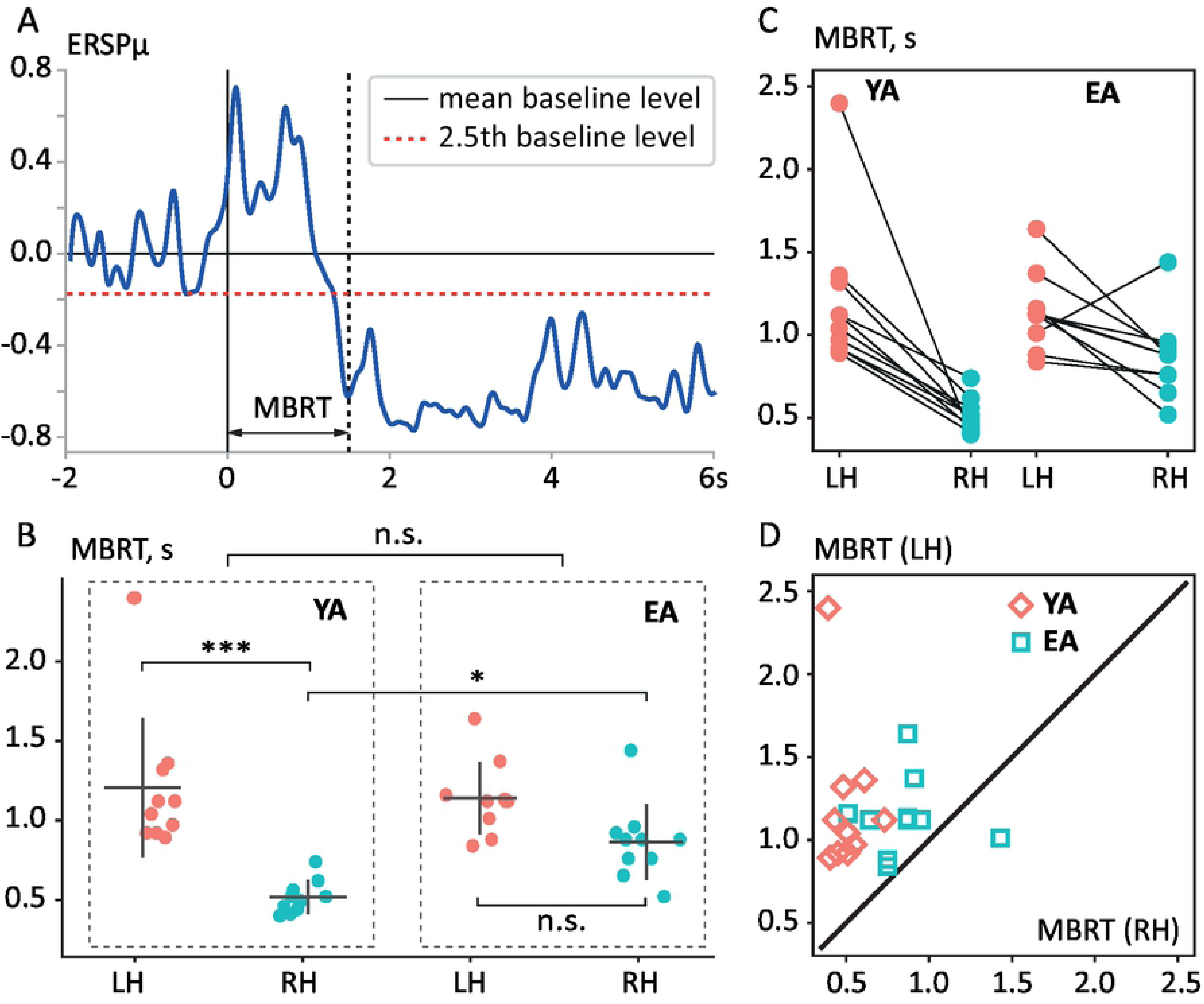
Motor brain response time. **A** An exemplary illustration of the MBRT estimation. The blue curve shows single-subject ERSP_*µ*_ at the C4 sensor averaged over 15 LH epochs. Black solid and red dashed horizontal lines indicate mean and 2.5^*th*^ percentile level of the baseline ERSP_*µ*_, respectively. Black solid and black dashed vertical lines show the beginning of the audio command and estimated motor brain response, respectively. **B** Distribution of MBRT across subjects in each (group,condition)-set. Here, ‘*’ indicates *p* < 0.05, ‘***’ indicates *p* < 0.001 and ‘n.s.’ stands for insignificant difference. **C** Stripcharts of linked observations MBRT(LH) and MBRT(RH) within each group. **D** Scatterplot of paired observations MBRT(LH) versus MBRT(RH) for each subject. Here, the diagonal line is MBRT(LH)=MBRT(RH), pink diamonds indicate subjects from YA group, and blue squares — subjects from EA group.

#### Within-subject time-frequency analyses

We performed within-subject spatio-temporal clustering analyses to reveal arrays of sensors associated with the motor-related brain activity for each age group and experimental condition. We considered baseline-corrected topo-maps averaged in the frequency bands of interest in the non-overlapping 0.2 s windows. Pairwise comparison of (time,sensor)-pairs was performed via one-sampled t-test (*DF* = 9, *p*_*pairwise*_ = 0.01, *t*_*critical*_ = ±2.821) and spatio-temporal clustering was assessed using non-parametric permutation test with *r* = 2000 random permutations (*p*_*cluster*_ = 0.01) following Maris and Oostenveld [33].

#### Between-subject time-frequency analyses

During between-subject analyses, we compared brain activity of the age groups in the same experimental conditions. Again, we considered baseline-corrected topomaps averaged in the frequency bands of interest in the non-overlapping 0.2 s windows. Effect of interest was evaluated at each (time,sensor)–pair using two-tailed *F* -test for independent samples (*DF*1 = 1, *DF*2 = 18, *p* = 0.05, *F*_*critical*_ = 5.978) and spatio-temporal clustering was assessed using non-parametric permutation test with *r* = 2000 random permutations (*p*_*cluster*_ = 0.05) [33].

#### Mixed-design analyses

Based on the results of within- and between-subject spatio-temporal clustering analysis, we localized the effect of significant spectral power change in the spatio-temporal domain. Further, for each (group, condition)-set we averaged spectral power over the corresponding spatio-temporal clusters and compared it using mixed-design ANOVA.

### Functional connectivity analysis

Functional connectivity measures the similarity of activation in the different brain regions based on the recorded signals of brain activity. According to the review papers [34, 35], there exists a variety of functional connectivity metrics that evaluate this similarity in the different aspects. Moreover, functional connectivity analysis based on EEG or MEG recordings suffers from such problems as volume conduction/field spread effect, signal-to-noise ratio, common input, etc. due to the nature of these neuroimaging techniques [36]. Thus, the choice of the particular functional connectivity measure requires both a prior knowledge about the analyzed neuronal processes and an understanding of possible problems that may potentially interfere with the adequate interpretation of functional connectivity results.

#### Functional connectivity measure

In accordance with the prior knowledge that motor-related activity is associated with certain frequency bands, first of all we expect the similarity of oscillatory behavior in remote brain regions in terms of phase-locking. Among the variety of FC measures based on the phase-synchronization, phase lag index (PLI) seems to be an appropriate metric [37]. PLI is robust to the common source problem as it ignores simultaneous phase similarity, less sensitive to the intrinsic EEG noise and allows reasonable interpretation of the obtained results. PLI is traditionally defined as:

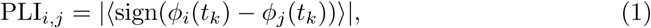

where *φ*_*i,j*_(*t*) are phases of signals at *i*^*th*^ and *j*^*th*^ EEG sensors introduced via Hilbert transform and operator 〈•〉 averaging over time points *k*. It clearly follows from Eq. (2), that PLI lies between 0 and 1, where PLI = 1 corresponds to perfect phase-locking and PLI = 0 implies a complete lack of synchrony.

*PLI* is also formulated in the frequency domain. In this case, the definition of PLI given in Eq. (2) is rewritten as:

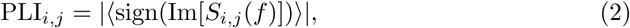

where *S*_*i,j*_ is a complex-valued Fourier-based cross-spectrum of *i*^*th*^ and *j*^*th*^ time-series and *f* covers the frequency band of interest. Frequency-domain definition of PLI is implemented in MNE package and has been used in this study to reveal motor-related functional connectivity.

#### Adjacency matrix

The functional connectivity structure in each of the frequency bands of interest was presented by symmetric adjacency matrix sized (31 × 31). For each of *n* = 20 participants we calculated *k* = 15 connectivity matrices in both experimental conditions (LH and RH) during the premotor interval 0 ÷ 1.25 s contrasted by the baseline connectivity (− 1.25 ÷ 0 s). Baseline contrast was applied to exclude false links, which could potentially arise due to the age-related changes in the resting-state functional connectivity. Then, for each subject, we computed mean connectivity matrices, averaged over *k* = 15 matrices, in both experimental conditions. To highlight statistically significant changes in functional connectivity related to the factor of age, we provided a between-subject analysis of the mean functional connectivity matrices. To address this issue we performed element-wise comparison of mean connectivity matrices for each type of movements between age groups using unpaired t-test with *p*_*pairwise*_ = 0.05 (*dF* = 9, *t*_*critical*_ = 2.101). Multiple comparison problem (MCP) was addressed via non-parametric permutation test with *r* = 2000 random permutations and *p*_*perm*_ = 0.05 [33].

## Results

### Motor brain response time analysis

First, we evaluated the effect of aging on the MBRT, i.e., the duration of the time interval required for the brain to activate a corresponding motor area for both groups. We estimated MBRT for each subject in both experimental conditions (Fig. 2B) and compared the results taking into account Age and Movement Type factors together (see Tables 1 and 2 in Section “Supplementary Materials”). The mixed-design ANOVA test revealed the significant effect of the Movement Type on the MBRT (*F* (1, 18) = 26.748, *p* < 0.001) while the factor of Age was insignificant (*F* (1, 18) = 2.626, *p* = 0.123). Post hoc comparison via paired t-test indicated that the mean MBRT for the LH condition (M=1.173, SD=0.341) was significantly higher than the RH condition (M=0.691, SD=0.248). Thus, in the case of the dominant hand movement (RH condition), the motor cortex activated faster in 19 of 20 subjects excluding 1 EA subject (Fig. 2C and D). Also, two EA subjects demonstrated almost close motor response times in both experimental conditions.

**Table 1.** Motor brain response time (Two-way mixed-design ANOVA Summary)

| Cases | DF1 | DF2 | Mean Square | F | p |
| --- | --- | --- | --- | --- | --- |
| Age (between-subject) | 1 | 18 | 0.199 | 2.626 | 0.123 |
| Movement Type (within-subject) | 1 | 18 | 2.325 | 26.748 | < .001*** |
| Movement Type * Age | 1 | 18 | 0.432 | 4.967 | 0.039* |

**Table 2.**
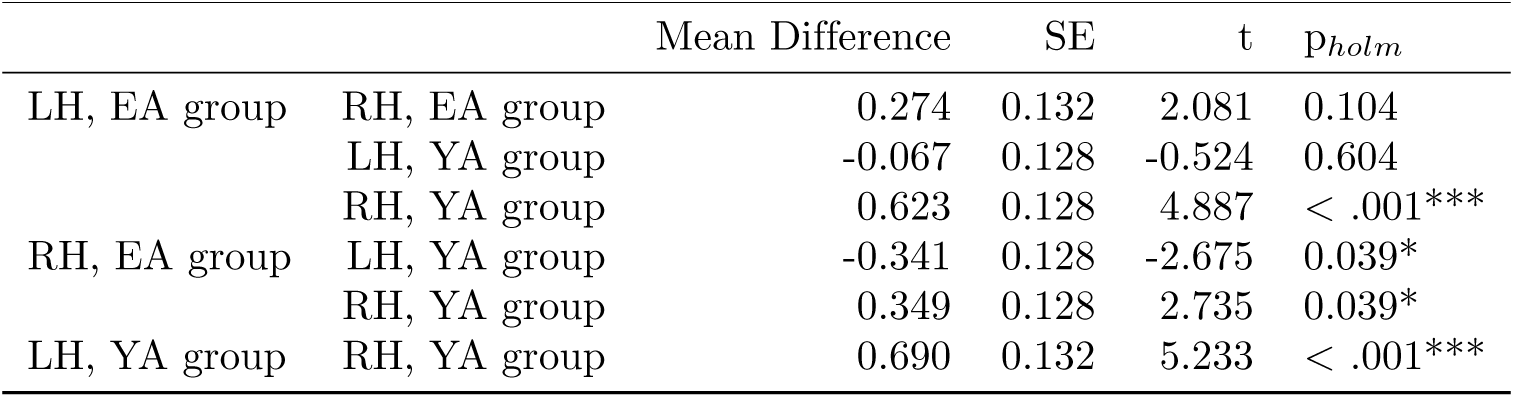
Motor brain response time (Post hoc comparisons)

Moreover, there was a significant interaction between the Movement Type and the Age of the participants (*F* (1, 18) = 4.967, *p* = 0.039). We could interpret this interaction as meaning that the Movement Type influenced MBRT differently in EA and YA groups. Particularly, MBRTs were similar in the LH condition (EA: M=1.139, SD=0.219; Young: M=1.206, SD=0.427), while YA group reacted significantly faster in RH condition (EA: M=0.865, SD=0.230; Young: M=0.516, SD=0.098).

### Within-subject time-frequency analysis

Based on the above MBRT analysis, we assumed that age-related changes affecting the speed of brain motor activation should be found in the pre-motor period. With this aim, we performed within-subject spatio-temporal clustering analysis of the spectral power in the theta and alpha/mu frequency bands for each (group, condition)-set in the premotor period (0 ÷ 1.2 s). Fig. 3 shows the results of within-subject clustering analysis in the LH condition for both groups of subjects. It is seen that in the LH condition (non-dominant hand movement), brain activation in both YA and EA groups proceeds similarly. Specifically, the suppression of mu-rhythm in the motor cortex at 0.8 ÷ 1.2 s related to the motor execution control was preceded by the theta-band activation during the period from 0.2 s to 0.8 s. In the YA group, pre-motor theta-band activation involved sensors in the motor, frontal, and bilateral temporal areas. In the EA group, strong theta-band synchronization spanned widely across the frontal and motor areas. Thus, in LH condition, both groups shared the same activation mechanism and timing of the motor initiation process.

**Fig 3.**
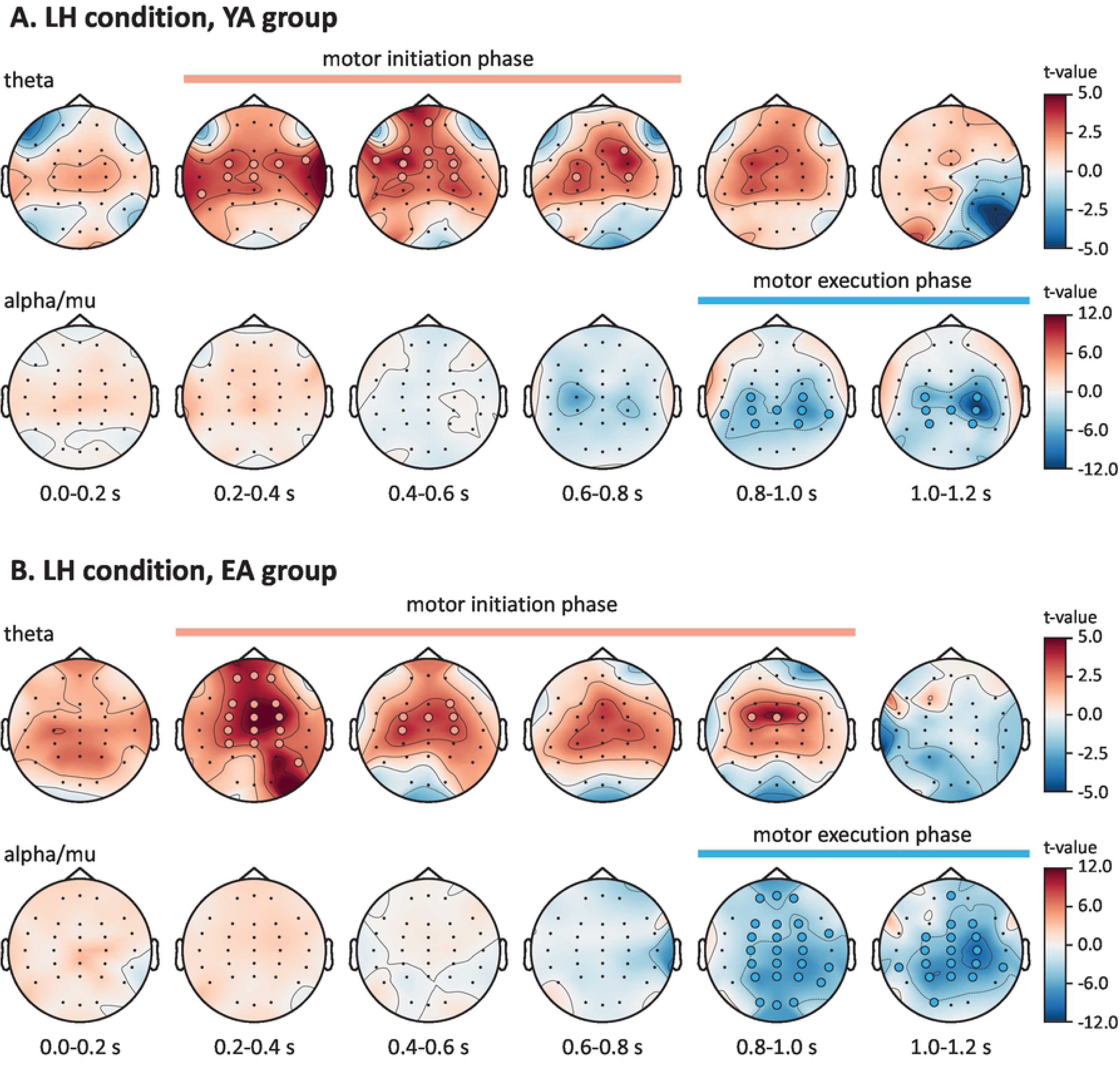
Sensor-level within-subject analyses of the pre-motor activity (LH condition). Baseline-corrected topomaps of the theta-band (upper row) and alpha/mu-band (lower row) spectral power for the YA (**A**) and EA (**B**) groups. Red and blue dots indicate clusters of significant spectral power increase and decrease respectively. Pairwise comparison is performed via t-test for related samples with *p*_*pairwise*_ = 0.01 (*dF* = 9, *t*_*critical*_ = ±2.821) and cluster-based analysis is performed via non-parametric permutation test with *p*_*cluster*_ = 0.01.

On the contrary, the way of cortical activation during the pre-motor period in the RH condition (dominant hand movement) was different in considered age groups (Fig. 4). In the YA group, the theta-band spectral power did not change significantly during the pre-motor period and the mu-band suppression in the motor cortex began earlier compared to the LH condition (0.6 ÷ 0.8 s instead of 0.8 ÷ 1.0 s). At the same time, in the EA group, the pre-motor brain dynamics in the RH condition completely reproduces the LH condition scenario in terms of spatio-temporal localization of synchronization and desynchronization in the theta and alpha/mu bands, respectively.

**Fig 4.**
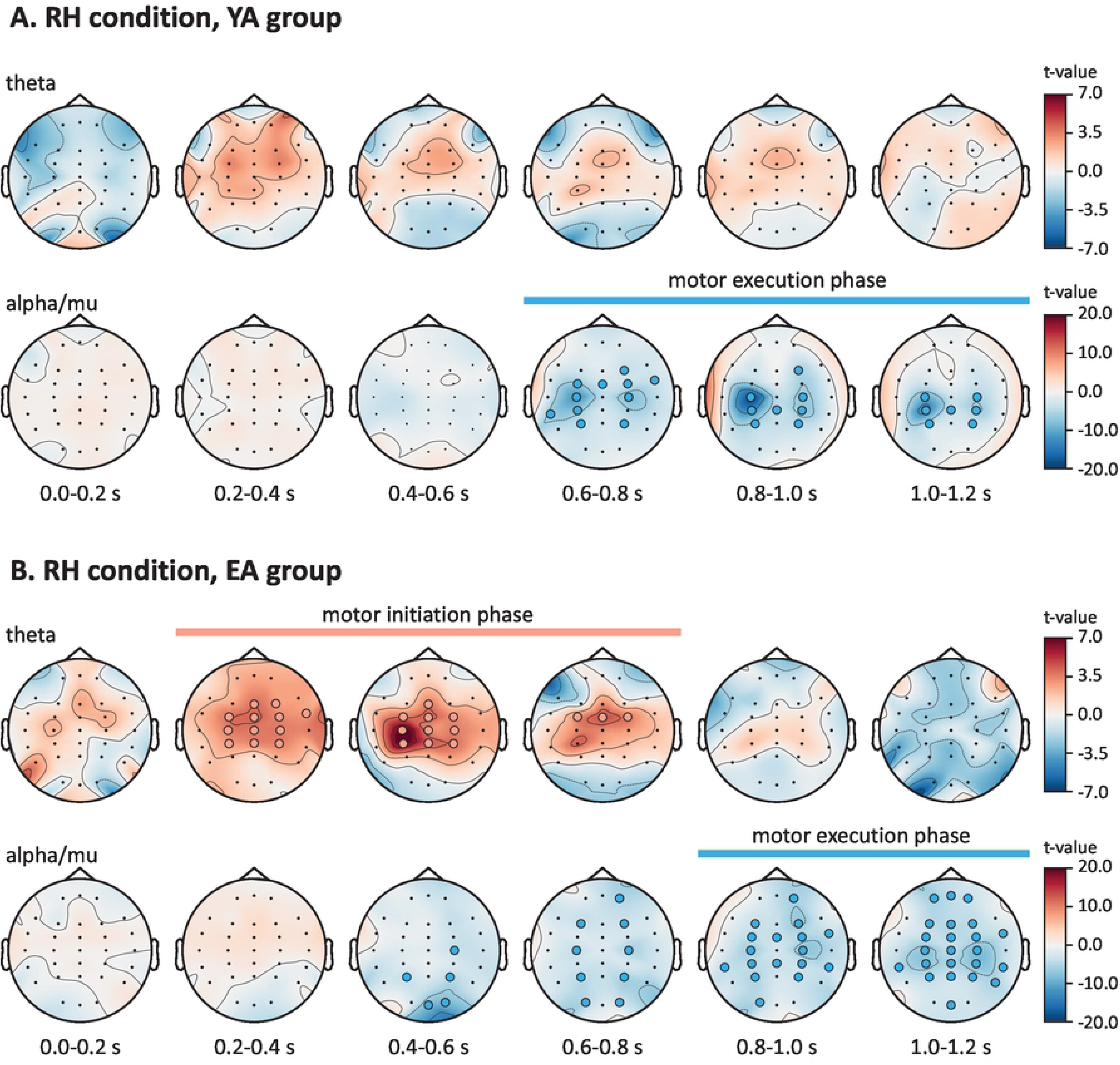
Sensor-level within-subject analyses of the pre-motor activity (RH condition). Baseline-corrected topomaps of the theta-band (upper row) and alpha/mu-band (lower row) spectral power for the YA (**A**) and EA (**B**) groups. Red and blue dots indicate clusters of significant spectral power increase and decrease respectively. Pairwise comparison is performed via t-test for related samples with *p*_*pairwise*_ = 0.01 (*dF* = 9, *t*_*critical*_ = ±2.821) and cluster-based analysis is performed via non-parametric permutation test with *p*_*cluster*_ = 0.01.

### Between-subject time-frequency analysis

To address the age-related changes of the pre-motor theta-band activation in detail, we provided a between-subject analysis of spectral power topo-maps. Fig. 5A shows the results of between-subject spatio-temporal clustering analysis performed separately in each experimental condition. In LH condition, the significant between-subject difference in the theta-band activation was not observed. On the contrary, the between-subject differences were found in the spatial cluster, which included Cp3, Cpz, and Cp4 sensors (dorsal stream region of the sensorimotor area) in a 0.4-0.6s window before the RH movement execution. Thus, we localized the effect in the spatio-temporal domain.

**Fig 5.**
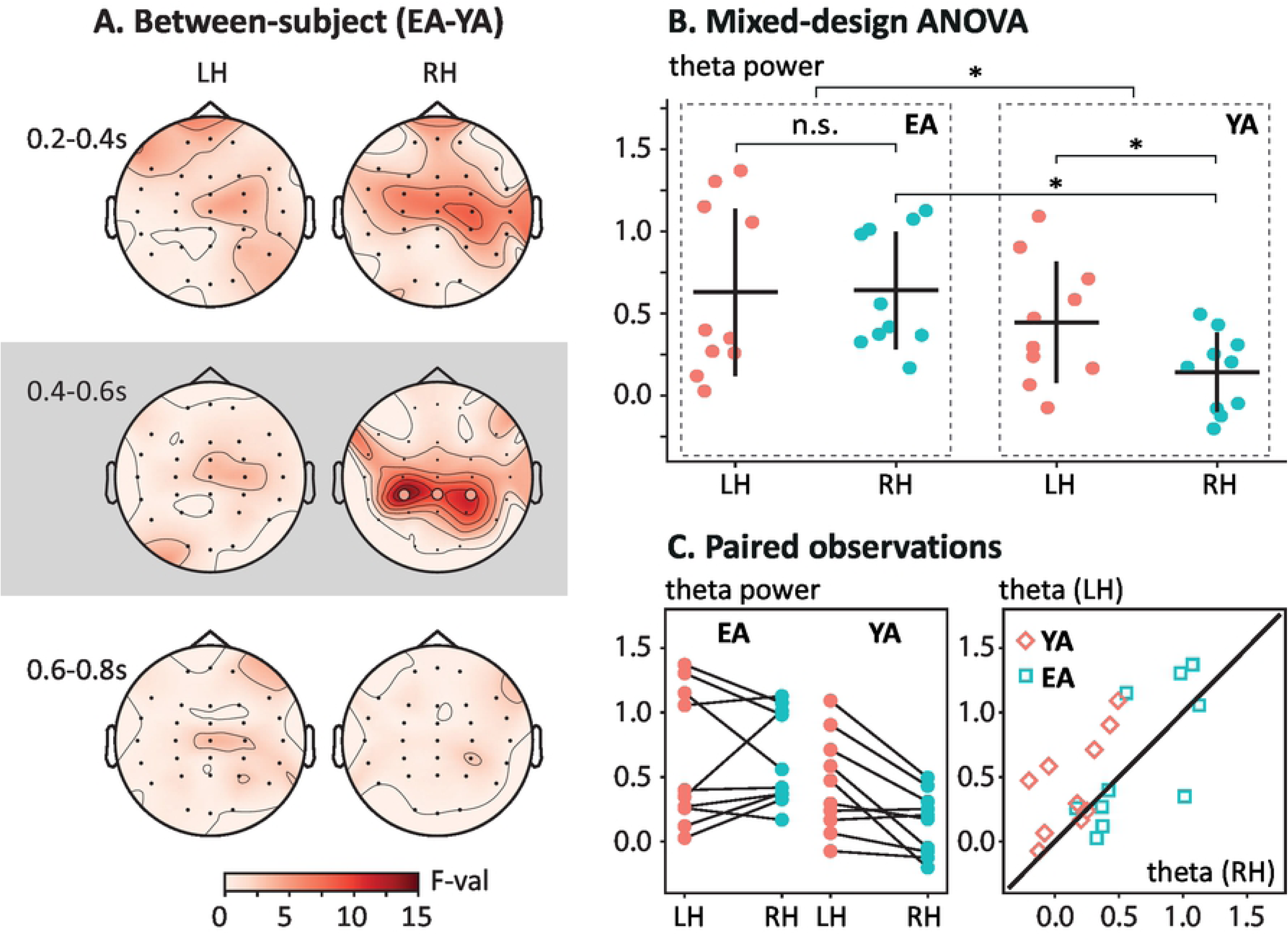
Analyses of the pre-motor theta-band activation. **A.** Between-subject analysis of the event-related theta-band spectral power preceding the LH (left column) and RH (right column) movements execution. Red circles indicate cluster of sensors with significant differences via non-parametric clustering test. Pairwise comparison is performed via two-tailed F-test for independent samples with *p*_*pairwise*_ = 0.05 (*dF*1 = 1 and *dF*2 = 18, *F*_*critical*_ = 5.978) and cluster-based analysis is performed via non-parametric permutation test with *p*_*cluster*_ = 0.05. **B.** Distribution of the event-related theta-band spectral power averaged over the sensors Cp3, Cpz, and Cp4 forming a positive cluster in **A** across subjects in each (group, condition)–set. Here, ‘*’ indicates *p* < 0.05 and ‘n.s.’ stands for insignificant difference. **C.** Strip-charts (left) and scatterplot (right) of paired observations within each group. Here, diamonds indicate subjects from YA group and squares indicate subjects from EA group.

To estimate age-related differences of theta-band activation taking into account both Age and Movement Type factors, we compared mean theta-band spectral power over the evaluated spatio-temporal cluster via mixed-designed ANOVA (Fig. 5B). The mixed-design ANOVA test revealed the significant effect of the Age on the pre-motor theta-band power (*F* (1, 18) = 4.636, *p* = 0.045), while the factor of Movement type was insignificant (*F* (1, 18) = 4.158, *p* = 0.056). Mixed-design ANOVA summary presented in detail in Tables 3. Post hoc comparison via unpaired t-test indicated that the mean pre-motor theta-band spectral power for the EA group (M=0.636, SD=0.429) was significantly higher than the YA group (M=0.294, SD=0.336). Also, similarly to the MBRT analysis, there was a significant interaction between the Movement Type and the Age of the participants (*F* (1, 18) = 4.770, *p* = 0.042). We interpret this interaction as follows: pre-motor theta-band power was similar for LH condition the EA and YA groups (EA LH: M=0.631, SD=0.524; YA LH: M=0.446, SD=0.376), while the YA group demonstrated higher pre-motor theta-band power in RH condition (EA RH: M=0.641, SD=0.365; YA RH: M=0.142, SD=0.242). Post hoc comparison results presented in detail in Table 4.

**Table 3.** Pre-motor theta-band spectral power (Two-way mixed-design ANOVA Summary)

| Cases | DF1 | DF2 | Mean Square | F | p |
| --- | --- | --- | --- | --- | --- |
| Age (between-subject) | 1 | 18 | 1.170 | 4.636 | 0.045* |
| Movement Type (within-subject) | 1 | 18 | 0.216 | 4.158 | 0.056 |
| Movement Type * Age | 1 | 18 | 0.247 | 4.770 | 0.042* |

**Table 4.**
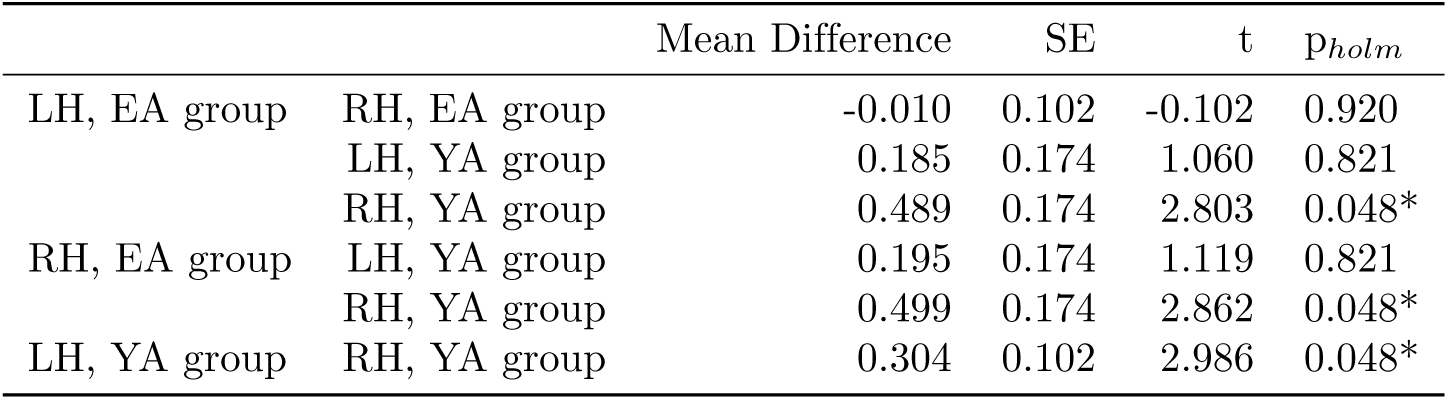
Pre-motor theta-band spectral power (Post hoc comparisons)

| | | Mean Difference | SE | t | $p_{holm}$ |
| --- | --- | --- | --- | --- | --- |
| LH, EA group | RH, EA group | -0.010 | 0.102 | -0.102 | 0.920 |
|  | LH, YA group | 0.185 | 0.174 | 1.060 | 0.821 |
|  | RH, YA group | 0.489 | 0.174 | 2.803 | 0.048* |
| RH, EA group | LH, YA group | 0.195 | 0.174 | 1.119 | 0.821 |
|  | RH, YA group | 0.499 | 0.174 | 2.862 | 0.048* |
| LH, YA group | RH, YA group | 0.304 | 0.102 | 2.986 | 0.048* |

### Functional connectivity analysis

To support and extend our observations of the pre-motor cortical activation, we explored age-related changes in terms of the underlying functional interactions between remote brain regions. As we demonstrated above that theta-band activity preceded the motor execution phase, we provided a between-subject comparison of the sensor-level theta-band functional connectivity matrices in the pre-motor stage of the RH condition (0 ÷ 1.25 s). As seen in Fig. 6A, the distributed frontoparietal network, including the midline connections, was strongly coupled in YA group compared to EA subjects. At the same time, we found the significant bilateral coupling increase between the motor cortex, temporal, and frontal lobes in EA participants (Fig. 6 B). Here, the large-scale neuronal communication was provided through the strong hub located in the primary motor cortex (Cz–sensor), which aggregated sensory information via coupling with the bilateral cortical sensorimotor circuits (C3–TP7 and C4–TP8), temporal area (Cz–FT7, Cz–T7) and transferred it for further processing to the frontal area (Cz–F3, Cz–F4, Cz–Fc4).

**Fig 6.**
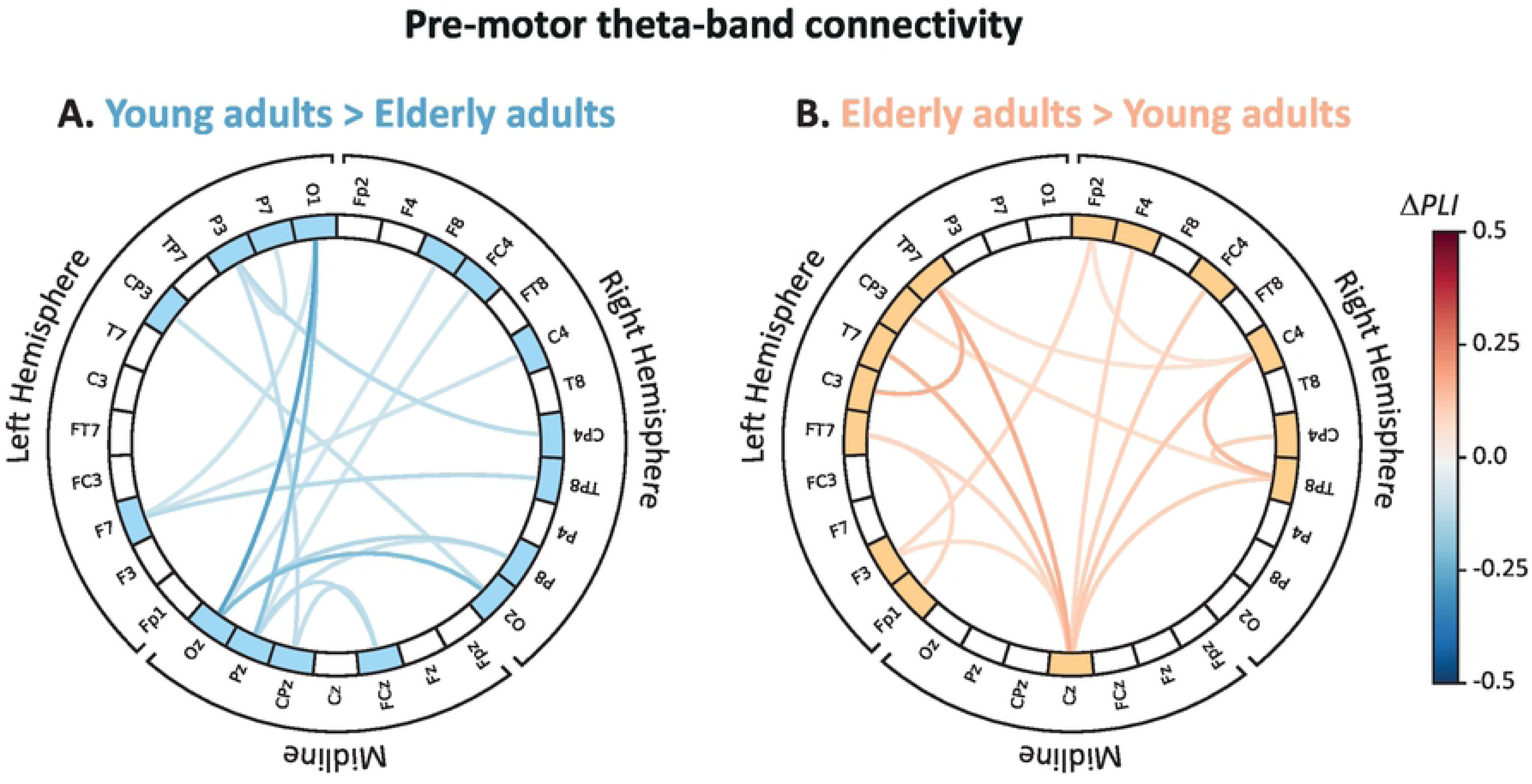
Between-subject analysis of the pre-motor theta-band functional connectivity in the RH condition. **A.** Significantly stronger coupling in YA compared to the EA. **B.** Significantly stronger coupling in EA compared to the YA. Here, ∆PLI defines the difference between group-level mean functional connectivity (EA versus YA). Element-wise comparison of mean connectivity matrices between Age groups was performed via unpaired t-test with *p*_*pairwise*_ = 0.05 (*dF* = 9, *t*_*critical*_ = ±2.101).

## Discussion

We considered the effect of healthy aging on the cortical activation in the motor initiation phase during the controlled execution of fine motor tasks – squeezing a hand into a fist paced by the audio command. We found that the time required for motor-related mu-band desynchronization, which we referred to as a motor brain response time (MBRT), was increased in the elderly subjects compared to the younger control group. Based on the results of time-frequency and functional connectivity analyses, we found that more extended motor response was preceded by the increased theta-band activation in the dorsal stream region of the sensorimotor cortex and stronger coupling between the primary motor cortex, bilateral temporal areas, and frontal lobe. Further, we discuss our results in the context of possible mechanisms supporting the motor initiation slowdown.

We observed the significant age-related differences in the MBRTs, which demonstrated a higher speed of the motor initiation in the case of the dominant (right) hand task in younger participants compared to the elderly adults. At the same time, motor initiation was equally slow during the non-dominant (left) hand task in both age groups. Moreover, MBRTs of elderly adults in both conditions approached the level of the non-dominant hand in younger subjects. Based on these findings, we suggested that the neuronal mechanisms supporting right-hand dominance are impaired under healthy aging. Despite the conflicting evidence in the literature, our results are consistent with several studies showing a similar effect. First, T. Kalisch et al. [8] demonstrated the behavioral decline in the dominant hand performance leading to ambidexterity in elderly adults. The authors argued that their findings could be explained by the mechanism of use-dependent plasticity [16], causing the degradation of well-trained motor functions due to the reduced activity and sedentary lifestyle of elderly individuals. Also, J. Langan et al. [11] supported these results and showed less-lateralized task-related motor activity in elderly adults compared to the younger control group. They found that longer reaction time in elderly adults was correlated with greater activation of the ipsilateral primary motor cortex during the motor task performance and weaker resting-state interhemispheric coupling, which was also observed in Refs. [10, 38, 39]. Described changes provided the compensatory mechanism to maintain the level of motor performance consisting in the reorganization of functional networks aimed at overcoming the age-related chemical and structural changes [14, 15]. Our results also evidence the motor-related over-activation of the brain areas in elderly adults as a large cluster of mu-band desynchronization covering additional areas of frontal, motor, parietal, and temporal regions (see Fig. 3 and Fig. 4).

However, the aforementioned mechanisms are not the only ones that support the brain’s motor response slowdown. Our results showed that the prevalent theta-band activation in the sensorimotor and frontal areas preceded the motor-related mu-band desynchronization during the non-dominant hand movements in both groups and the dominant hand movements in elderly adults. Mechanism of motor initiation related to the increased theta activity is explained by the Bland’s sensorimotor integration model. In their early works on rodents [40, 41], B.H. Bland with colleagues treated the hippocampal formation theta activity as a communication channel between the sensory processing and movement initiation. Further, the Bland’s model was extended to the human brain in a series of works by J.B. Caplan et al. [19, 42]. In their studies, they concluded that while mu-band suppression (a traditional hallmark of the motor-related brain activity) reflected cortical activation directly during the motor task execution, the increased theta power between stimuli presentation and motor execution was associated with sensorimotor integration similarly to rodents. Along with this, several EEG-studies reported the increase of theta-band power during the planning phase in the choice-related, catching, and imagery motor tasks [43–45]. Specifically, M. Tambini et al. [44] demonstrated a positive correlation between theta-power and task performance. On the contrary, we found that increased theta-band power was associated with prolonged motor initiation. It should be noted that the significant increase of the theta-band power related to the dominant hand decline in elderly adults was observed in the dorsal stream region of the sensorimotor cortex and associated with the motor planning but not with audio command processing. Following the recent study by J. Dushanova et al. [46], such a result should be explained by the different strategies of the motor task initiation between age groups. While the degraded plasticity in elderly adults requires higher cortical activation for motor planning, younger subjects optimize their cognitive resources for the familiar and well-trained motor task accomplishment. The latter was represented as a lower theta-band activation. Therefore, less effective use of cognitive resources prolonged the motor planning phase in elderly adults compared to the younger control group during the dominant hand tasks.

These conclusions about the age-related differences in the motor initiation strategies during the dominant hand activity were supported and extended by the pre-motor functional connectivity analysis. The differences in theta-band functional connectivity pattern between two groups could be interpreted as a meaning of the different mechanisms of motor planning in elderly adults and young subjects. First, we showed the stronger neuronal interaction within the frontoparietal cognitive network involving midline connections in younger adults compared to the elderly group. It reflects the increased perceptual-motor facility and motor working memory [47–50]. We suppose that in young adults, initiation of the familiar motor activity emphasizes motor working memory and enables the formation and processing of the motor memories, i.e., the stored information about the motor action obtained from prior experience, for accurate motor performance [51]. The process of memory representation is fast and efficient in terms of cognitive demands. On the contrary, in elderly adults, functional connectivity inferred stronger coupling between the frontal, motor, and bilateral temporal areas with a central hub in the primary motor cortex. As the working memory decline with age is well-documented [52–54], we suggest that memory representation of motor actions is less accessible in elderly adults. Thus, the involvement of sensorimotor integration mechanisms relative to the Bland’s Type 1 motor-related theta activation [19], which requires higher cognitive resources, is prevalent in the elderly group. Based on the above functional connectivity results, we conclude that the difference of the motor planning mechanisms is more demanding and could not be optimized as well as in younger subjects causing the significantly delayed motor initiation.

## Conclusion

Elderly adults exhibited the approach to ambidexterity in term of the slowdown in cortical activation related to the execution of the dominant hand task. We demonstrated that the observed age-related loss of the dominant hand advance was due to the difference in motor initiation strategy between elderly adults and young subjects. Our results suggest that while young participants tend to activate motor working memory to initiate the dominant hand movement, elderly subjects involve a more demanding process of sensorimotor integration similar to the Bland’s Type 1 motor-related theta activation, which significantly reduces the speed of motor planning.

## Author Contributions

Conceptualization: N.F., V.M. and A.H.; methodology: N.F.; data curation: V.G. and A.K.; software: N.F. and E.P.; formal analysis: N.F., E.P. and V.M.; investigation: N.F., E.P., Z.W.; writing–original draft preparation: N.F., E.P. and A.H.; writing–review and editing: N.F., V.M. and A.H.; visualization: N.F.; supervision: N.F. and A.H.; project administration: N.F. and A.H.; funding acquisition: N.F. and A.H. All authors have read and agreed to the published version of the manuscript.

